# *Txn1* mutation is a monogenic cause of chronic kidney disease associated with mitochondrial dysfunction in rats

**DOI:** 10.1101/2023.08.14.553187

**Authors:** Iori Ohmori, Mamoru Ouchida, Yoshiko Hada, Haruhito A. Uchida, Shinya Toyokuni, Tomoji Mashimo

## Abstract

Oxidative stress plays a significant role in the progression of chronic kidney disease. Thioredoxin 1 (Txn1) is one of the enzymatic antioxidants to regulate redox balance. However, the molecular mechanisms by Txn1 affects renal homeostasis remain unclear. This study aimed at elucidating the pathophysiology of *Txn1* mutations in renal function. We used rats with the *Txn1-*F54L mutation generated by N-ethyl-N-nitrosourea mutagenesis. Laboratory tests and pathological examinations were performed in wild-type (WT) rats and in rats with heterozygous and homozygous *Txn1*-F54L mutations. We performed RNA-seq analysis of the WT and homozygotes. To confirm phenotypic reproducibility, the *Txn1*-F54L mutation was generated in rats with different genetic backgrounds using CRISPR/Cas9 genome editing technology. *Txn1*-F54L mutant rats exhibited progressive albuminuria, hypoalbuminemia, hypercholesterolemia, and hypertension. Renal pathology revealed marked nephrosclerosis, tubular dilatation, interstitial fibrosis, and decreased number of mitochondria, mainly in the paroxysmal tubules. We confirmed a similar phenotype of chronic kidney disease (CKD) in different rat strains. RNA-seq showed the downregulation of mitochondria-related genes and significant upregulation of genes associated with inflammation, pyroptosis, apoptosis, and necroptosis in mutant rats. Our results show that the *Txn1* mutation is a monogenic cause of CKD termination. The underlying pathology involves several regulated cell-death pathways. Thus, our study provides a new animal model of oxidative stress-induced CKD. *Txn1-*F54L mutant rats will aid in developing therapeutic strategies for CKD.

**Translational Statement:** We found that the deficiency of thioredoxin (Txn1), which regulates oxidative stress, spontaneously caused chronic kidney disease (CKD) in rats. The Txn1-F54L (*Adem*) rat is a new model of CKD with complications such as anemia, hypertension, and cardiovascular disease. Renal pathology revealed nephrosclerosis, interstitial fibrosis, and mitochondrial damage. The molecular basis of the underlying pathologies included inflammation, pyroptosis, apoptosis, and necroptosis. These pathological changes are partially linked to renal diseases such as diabetic nephropathy, hypertensive nephrosclerosis, and ischemic reperfusion injury. *Adem* rats could help understand the common pathological mechanisms of these renal diseases and develop therapeutic strategies.

## INTRODUCTION

Chronic kidney disease (CKD) is caused by a combination of genetic and environmental factors. ^1^ However, the etiology differs between children and adults. Congenital anomalies of the kidney and urinary tract are the most frequent causes in children, ^2^ and diabetes and hypertension are the leading causes of CKD in adults. ^3^ More than 450 monogenic genes associated with CKD have been identified. ^1^ With advances in genetic analysis techniques, causative genes have been detected in approximately 30% of pediatric cohorts ^4^ and 5–30% of adult cohorts. ^5, 6^

In adults, diabetes, hypertension, and hyperlipidemia are often involved in the development and progression of CKD, and the complications of CKD exacerbate these conditions. Oxidative stress is involved in diabetes, hypertension, and hyperlipidemia. ^7, 8^ Thioredoxin (Txn1) is one of the enzymatic antioxidants such as superoxide dismutase, catalase, and glutathione peroxidase. Txn1 scavenges H_2_O_2_ (reactive oxygen species) and reduces oxidized proteins to maintain cellular redox balance. ^9^ Some studies have described a relationship between diabetic nephropathy and Txn1/thioredoxin-interacting protein (TXNIP) system. TXNIP plays a role in inhibiting Txn1, ^10, 11^ and high glucose levels upregulate its expression in humans and mice. ^12^ Overexpression of Txnip leads to apoptosis in the pancreatic beta cells, ^12, 13^ and Txnip knockdown prevents glucose-induced apoptosis in beta cells. ^14^ High levels of TXNIP have been observed in patients with diabetic nephropathy. ^15^ The evidence for the Txn1/TXNIP system being an aggravating factor in diabetes and diabetic nephropathy has been accumulating but is not unequivocal. Although TXNIP expression is elevated in the muscles of individuals with prediabetes and diabetes, no correlation between TXNIP SNP genotypes and type 2 diabetes has been found. ^16^ Epigenome-wide association studies have identified a significant association between type 2 diabetes and hypomethylation of TXNIP. ^17, 18^ Hypomethylation of TXNIP has been interpreted as an epigenetic consequence of hyperglycemic exposure rather than a cause of CKD. In animal research, *Txnip*-overexpressing mice exhibited enhanced severity of streptozotocin-induced diabetes but not kidney disease. ^19^ The precise molecular mechanisms underlying the Txn1/Txnip system in CKD remain largely unknown as the Txn1 KO mouse is embryonically lethal. ^20^ To our knowledge, there have been no reports of mutations in oxidative stress-related genes directly causing CKD in humans or animals.

Recently, we characterized an epileptic rat with a Txn1-F54L mutation. ^21^ Recombinant Txn1-F54L reduced insulin-reducing activity of Txn1 to approximately 1/3 of the wild type (WT). The rats exhibit age-dependent midbrain degeneration, elevated levels of H_2_O_2_ in the brain lesions, altered mitochondrial morphology of neurons, and higher susceptibility of fibroblasts from homozygous rats to apoptosis under oxidative stress. We named these rats *Adem* (Age-dependent mitochondrial cytopathy). ^22^ This study aimed to clarify whether the *Txn1* mutation causes CKD and its underlying pathology in this strain.

## METHODS

Detailed methods are provided in Supplementary Methods.

### Animals

All animal husbandry procedures were performed in accordance with protocols approved by the Institutional Experimental Animal Use Committee of Okayama University. Both sexes were included in the study.

Two rat strains were used in the present study. One group was F344/Nlc rats with *Txn1*-F54L mutation generated by N-ethyl-N-nitrosourea (ENU) mutagenesis, as previously reported. ^22^ The other was F344/Jcl rats with *Txn1*-F54L mutation generated by CRISPR/Cas9 genome editing ^22^ to confirm the reproducibility of the phenotype.

In this study, we monitored the general condition of rats for an extended period after epilepsy.

### Observation of general health and chemical tests

From 1 to 11 months of age, we measured body weight (laboratory electronic balance; A & D Company, Tokyo, Japan) and conducted chemical tests (Fuji DryChem7000V; Fuji Film, Tokyo, Japan) of blood and urine every two months for WT, heterozygous, and homozygous rats to check renal and liver function. The rats were weighed at 3.5 months, as homozygotes began to lose weight rapidly at approximately four months.

### Measurement of blood pressure

Blood pressure was measured using the non-invasive tail cuff method (BP-98AL; Softron Inc., Tokyo, Japan). The rats were moved to the laboratory room 1 h before the examination. Blood pressure was measured thrice once the rats calmed down. The average value was used as the measured value. Blood pressure was recorded every two months from the ages of 3 to 11 months.

### Histological assessment of the multiple organs

When the rats lost more than 20% of their body weight or their movement became sluggish in their home cage, they were euthanized and dissected (male heterozygotes: n = 9, female heterozygotes: n = 11, male homozygotes: n = 8, female homozygotes: n = 5). The primary body organs, including the heart, lungs, thymus, thoracic aorta, liver, kidneys, pancreas, and spleen, were examined. WT rats of the same age were used as controls (male: n = 8, female: n = 8). Each organ of each dissected rat was stained with hematoxylin and eosin (HE) stain. Kidneys were stained with periodic acid-Schiff (PAS), periodic acid methenamine silver (PAM), and Masson’s trichrome (MT), and blood vessels were stained with Elastica van Gieson (EVG).

### Immunohistochemistry analyses

A standard immunohistochemistry (IHC) protocol for 3,3′-Diaminobenzidine (DAB) staining was used to stain the kidneys of WT (male: n = 3, female: n = 3) and homozygotes (male: n = 3, female: n = 3) as previously reported. ^21^ The following primary antibodies were used: anti-8-OHdG for oxidative DNA damage, anti-NLRP3 and anti-GSDM for pyroptosis, anti-CASP3 for apoptosis, and anti-RIPK3 for necroptosis. The slides were counterstained with hematoxylin, mounted with dibutyl phthalate xylene (DPX), and observed using a BZX-700 multifunctional microscope (Keyence Corp., Osaka, Japan).

### Transmission electron microscopy

Kidney samples from WT (male: n = 2, female: n = 2) and homozygous (male: n = 2, female: n = 2) rats were immersed in 2% glutaraldehyde and 2% paraformaldehyde in 100 mmol/l phosphate buffer (PB) for 24 h at 4 °C. They were then postfixed with 2% osmium tetroxide in 100 mmol/l PB for 1.5 h at 4 °C, after which they were rinsed with 100 mmol/l PB and dehydrated through a graded series of ethanol treatments. The sections were embedded in Spurr resin (Polysciences Inc., Warrington, PA, USA), cut into ultrathin sections, and stained with uranyl acetate and lead citrate. The ultrathin sections were observed under a Hitachi H-7650 transmission electron microscope (TEM) (Hitachi High-Tech Corp., Tokyo, Japan). The length of mitochondria was measured by the National Institutes of Health (NIH) *ImageJ* software.

### RNA sequencing

RNA-seq analysis was performed on WT mice (n = 4) and homozygotes (n = 4) at 15 weeks of age. RNA was extracted from the renal cortex using a standard procedure (RNeasy Plus kit; Qiagen, Venlo, Netherlands). We used transcripts per million (TPM) values, which were normalized to the raw read counts for principal component analysis (PCA), hierarchical clustering, heatmapping, differential expression analysis, and gene ontology (GO) enrichment analysis.

### Confirmation of CKD reproducibility

We examined the CKD phenotype in CRISPR-Jcl wild-type (male: n = 3, female: n = 3) and homozygotes (male: n = 3, female: n = 3). Blood samples were collected from the tail vein of mice at 5, 10, and 15 weeks of age to measure blood urea nitrogen (BUN), total cholesterol (T-Cho), and albumin (Alb). When 20% weight loss or inactivation was observed, the rats were deeply anesthetized with isoflurane, followed by the sampling of whole blood and the main organs (heart, lung, thymus, thoracic artery, kidney, pancreas, liver, and spleen) (male: n = 2, female: n = 2). Each organ was subjected to HE staining. End-stage blood tests included BUN, T-Cho, Alb, phosphate (P), calcium (Ca), aspartate aminotransferase (AST), and creatine kinase (CK).

### Statistical analysis

Survival was evaluated using the Kaplan-Meier survival curves and compared between the different genotypes using log-rank test. Survival and chemical data were analyzed using EZR software. ^23^ Welch’s t-test was used for comparison between two groups. One-way analysis of variance (ANOVA) was used to compare the three groups. Statistical significance was set at *P* < 0.05. Bonferroni adjustment was used to determine statistical differences between the three groups. Results are expressed as the mean ± SEM.

## RESULTS

### *Adem* rats exhibit short lifespans, nephrosclerosis, and thymus involution

Homozygous and heterozygous rats lost weight after 3 and 11 months, respectively (Figures 1b and 1c). Survival curves were examined because some rats were debilitated and died. The median survival times of male homozygotes, female homozygotes, male heterozygotes, and female heterozygotes were 110 (95% confidence interval [CI] = 105-117), 119 (95% CI = 105-124), 303 (95% CI = 246-343), and 346 (95% CI = 264-398) days, respectively. In heterozygotes, the median survival time was significantly shorter in males than in females (*P* < 0.01) (Figure 1a). The faces of homozygotes and heterozygotes became edematous in the terminal state (Figures 1d and 1e). All euthanized rats were dissected to verify the cause of the illness. All mutant rats that underwent autopsy showed pale and swollen kidneys in homozygotes (Figure 1f) and hard and small kidneys in heterozygotes (Figure 1g). In addition, all mutant individuals showed marked regression of the thymus compared to WT individuals at the same age (Figures 1h and 1i).

**Figure 1.**
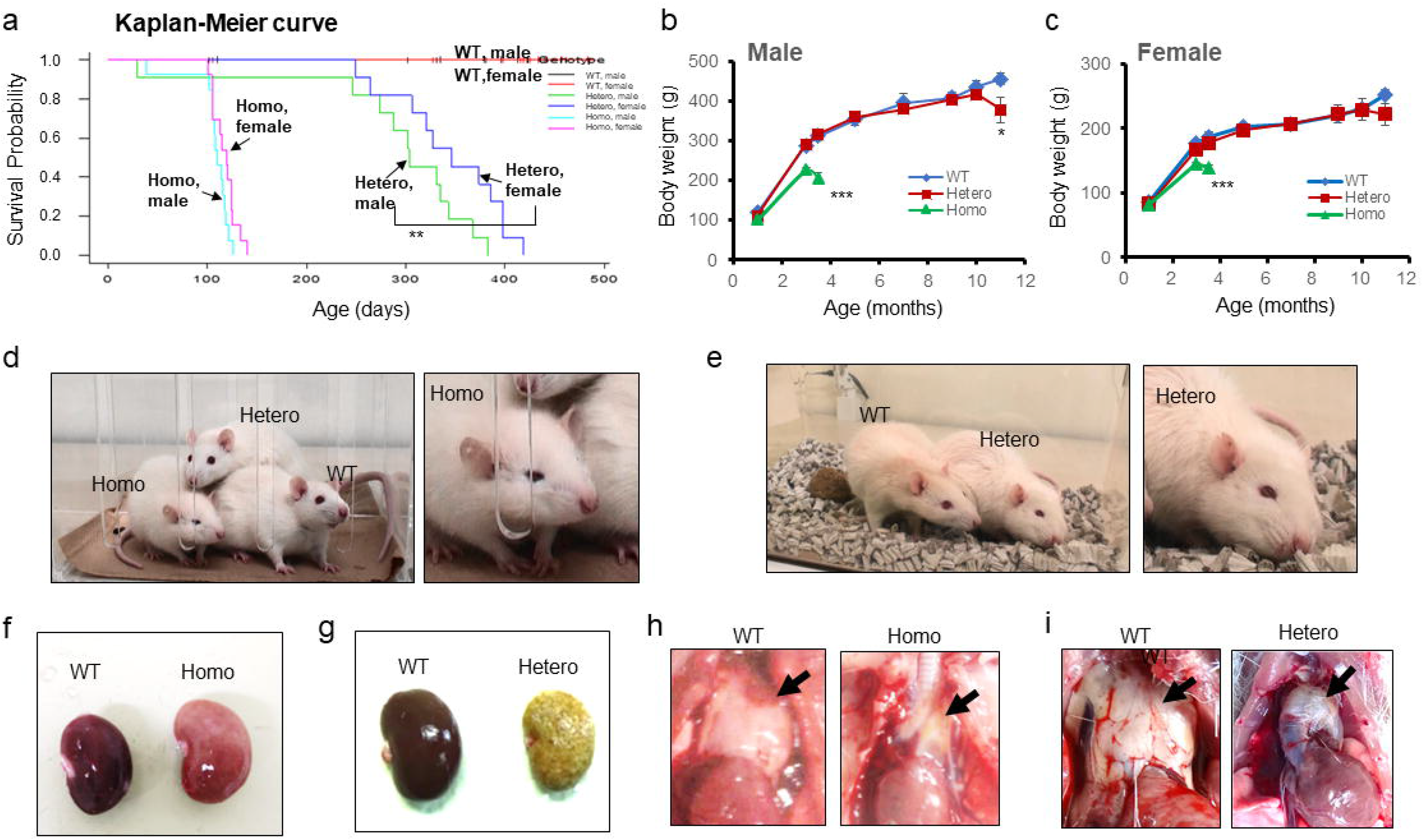
*Adem* rats exhibit premature death associated with renal disease and thymus involution. (**a)** Kaplan-Meier survival curves for each genotype and sex (male WT: n = 15, female WT: n = 12, male heterozygotes: n = 13, female heterozygotes: n = 13, male homozygotes: n = 11, female homozygotes: n = 11). (**b)** Male growth curve (WT: n = 12, heterozygotes: n = 6, homozygotes: n = 6). (**c)** Female growth curve (WT: n = 10, heterozygotes: n = 9, homozygotes: n = 9). **(d)** Representative appearance of the rats at four months showed facial edema in a homozygote. (**e**) Representative appearance of the rats at 11 months showed weight loss and edematous face. (**f**) Autopsy revealed that kidneys got pale and enlarged in homozygote. (**g**) Representative kidneys of heterozygote showed white and hardened. The thymus was present in the WT, whereas it was almost absent in a homozygote (**h**) and heterozygote (**i**). Black arrows indicate the thymus (WT) and thymus involution (mutants). * *P* < 0.05, *** *P* < 0.001

### *Adem* rats have CKD terminate in end-stage kidney disease (ESKD) without hyperglycemia

At the end stage, both heterozygotes and homozygotes showed a marked elevation in BUN (Figure 2a), notable decrease in serum albumin levels (Figure 2b), and hypercholesterolemia (Figure 2c). The level of AST, a marker of liver function, was not significantly elevated at any time point (Figure 2d). Hyperphosphatemia (Figure 2e) was also observed in homozygotes (*P* < 0.01); however, no significant difference in calcium levels was observed (Figure 2f). ALP levels were decreased in homozygotes (Figure 2g, *P* <0.01). Anemia was present at the terminus in both heterozygotes and homozygotes (Figure 2h). No hyperglycemia was observed in the mutants at any time (Figure 2i). HbA1C levels were measured at three months in homozygotes and 11 months in heterozygotes, with no significant differences (Figure 2h). Urinary albumin was gradually elevated in both heterozygotes and homozygotes (Figure 2k) (*P* < 0.01). Qualitative urinary test exhibited occult blood positive at the terminal stage (Figure 2l) while no urinary sugar at any points in both heterozygotes and homozygotes. Progression of CKD in homozygotes was dramatically faster than that in heterozygotes.

**Figure 2.**
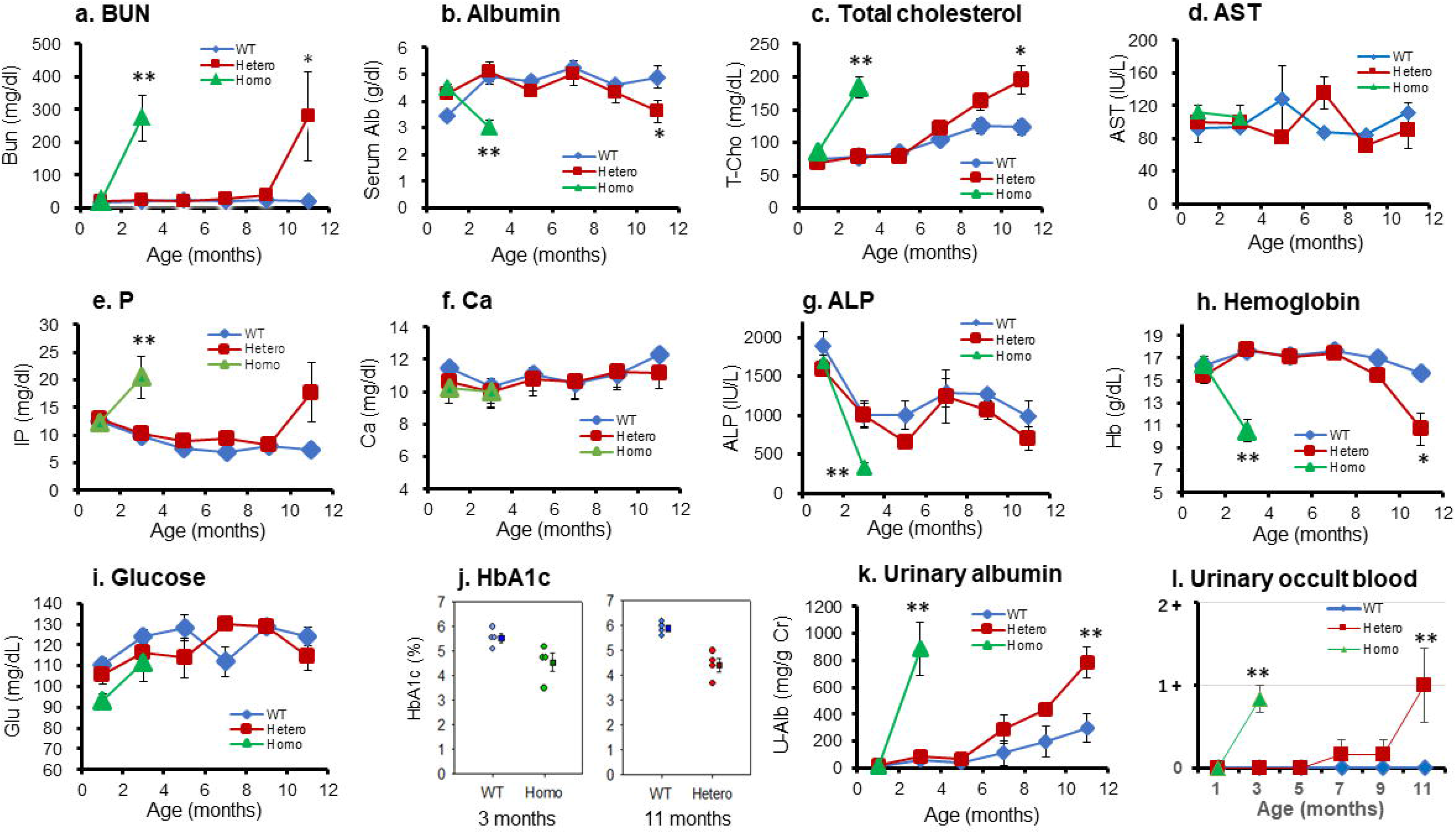
*Adem* rats develop CKD that eventually leads to renal failure. Chemical tests (male WT: n = 4, female WT: n = 4, male heterozygotes: n = 6, female heterozygotes: n = 19, male homozygotes: n = 6, female homozygotes: n = 9) include blood urea nitrogen (BUN) (**a**), serum albumin (**b**), total cholesterol (**c**), aspartate aminotransferase (AST) (**d**), serum phosphate (**e**), serum calcium (**f**), Alkaline Phosphatase (ALP) (**g**), hemoglobin (**h**), glucose (**i**), Hemoglobin A 1c (HbA1c) (**j**), urinary albumin (**k**), and urinary occult blood (**i**). * *P* < 0.05, ** *P* < 0.01

### Pathology of CKD

Both homozygous and heterozygous rats exhibited marked tubular dilatation. Renal lesions were extensive throughout the renal cortex. Nephrosclerosis and interstitial fibrosis were prominent in both the mutant genotypes (Figure 3a). Glomerular hyalinosis, mesangial expansion, crescents, and segmental and global sclerosis were also observed. The pathologies of the homozygotes and heterozygotes were similar. However, the heterozygous lesions progressed more slowly than the homozygous lesions and had more pronounced fibrosis. The TME in homozygous rats showed the fusion of glomerular foot processes, heterogeneous thickening, edema of the glomerular basement membrane, and nephrosclerotic lesions (Figure 3b).

**Figure 3.**
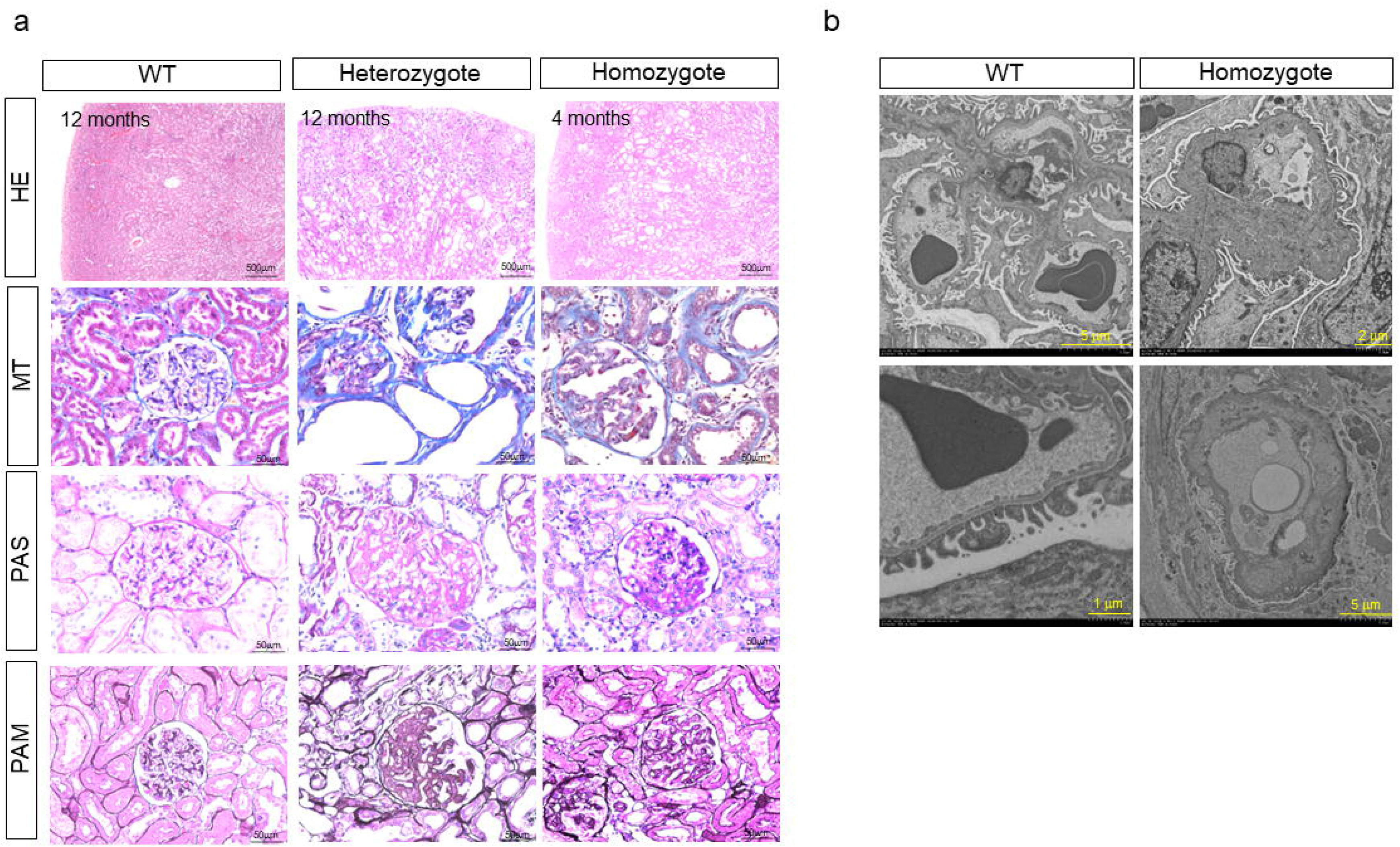
Pathology of chronic kidney disease. Representative HE, MT, PAS, and PAM stains are shown for the end-stage of homozygote (4 months), heterozygote (12 months), and healthy WT (12 months) (**a**). Both homozygous and heterozygous rats exhibited marked dilatation of tubules. Renal lesions were extensive in the renal cortex. Focal and global glomerulosclerosis, mesangial cell proliferation, hyalinosis, and interstitial fibrosis were prominent in both mutant genotypes. Representative TME findings for homozygotes (4 months) and WT (4 months) (**b**). Fusion of glomerular foot processes, heterogeneous thickening and edema of the glomerular basement membrane, and glomerulosclerotic lesions are observed in homozygous rats.

### Comorbidity of cardiovascular disease

Because of the close relationship between CKD, hypertension, and heart disease, we examined blood pressure and cardiovascular pathology. Hypertension appeared at three months in homozygotes and after seven months in heterozygotes compared to that in the WT (Figure 4a). The thoracic aorta was dilated in both homozygotes and heterozygotes (Figure 4b), and irregularities and tears in the vascular tunica media with calcification were observed (Figure 4c). These changes were compatible with Mönckeberg medial sclerosis. There were no atherosclerotic changes such as the accumulation of lipid deposits or inflammatory cell debris in the intimal architecture. In the myocardium, compared to the WT, homozygotes and heterozygotes showed lymphocytic infiltration, fibrosis (Figure 4e), and dilatation of intramyocardial vessels (Figures 4f and 4g). The degree of lymphocytic infiltration, fibrosis, and intramyocardial vasodilation was more pronounced in the heterozygotes than those in the homozygotes.

**Figure 4.**
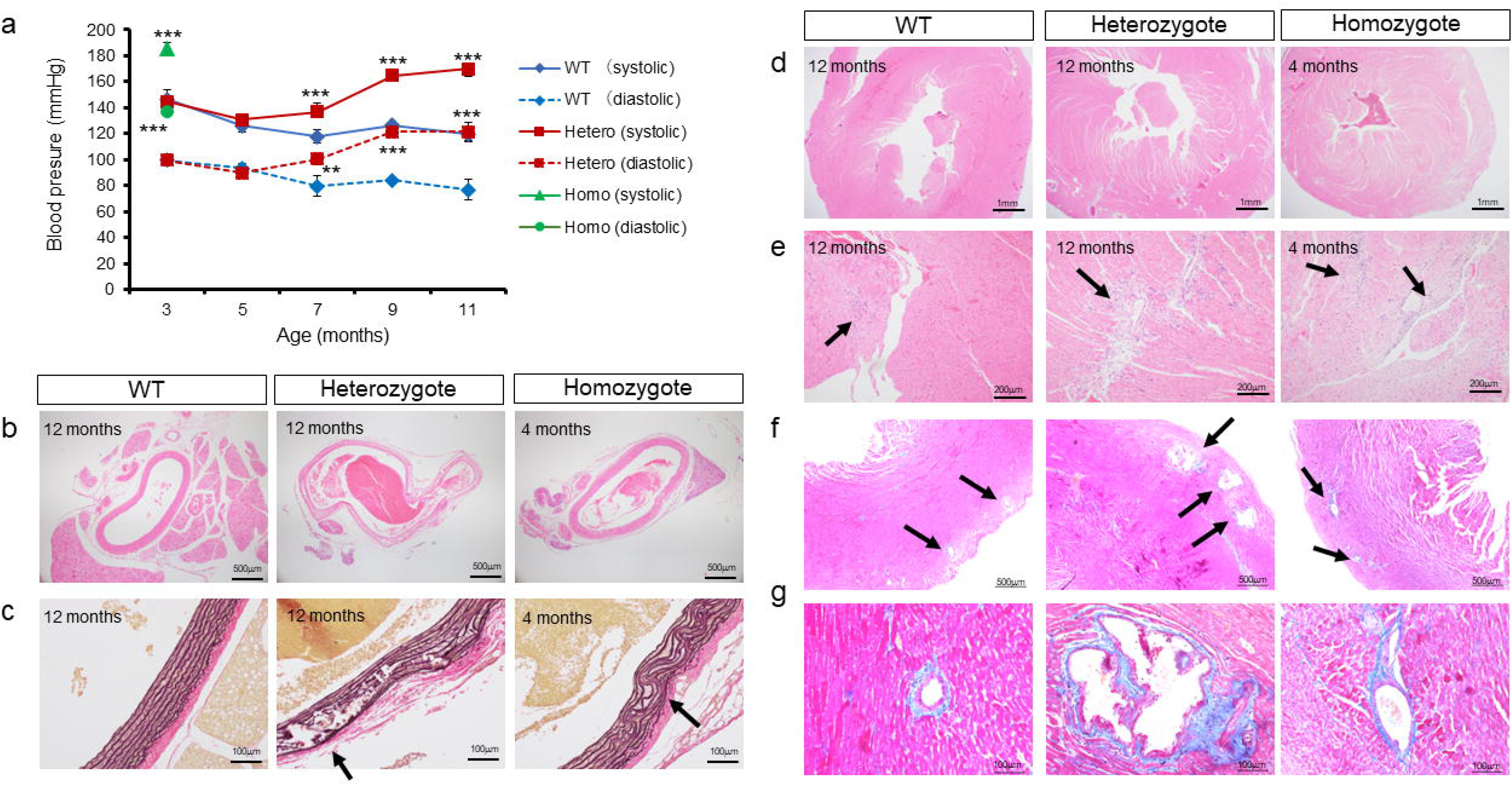
Comorbidity of hypertension and cardiovascular disease. Course of blood pressure for WT (n = 7), heterozygotes (n = 7), and homozygotes (n = 6) (**a**). Representative pathological observations in the end-stage of homozygotes (4 months), end-stage of heterozygotes (12 months), and healthy WT (12 months) are shown from (**b**) to (**g**). HE stain of thoracic aorta (**b**). Elastic van Gieson stain of thoracic aorta (**c**). HE stain of the hearts (**d** and **e**). MT stain of the hearts (**f** and **g**). ** *P* < 0.01, *** *P* < 0.001

Regression of the thymus and decreased lymphocytes in the spleen were also observed in other organs (Supplementary Figure S1). No significant changes were observed in the liver or islets of Langerhans in the pancreas. Dilated alveoli were observed in the lungs of some heterozygous individuals. There were individual differences in the lung findings.

### Downregulation of mitochondria-related genes and mitochondrial damage in the paroxysmal tubules

To elucidate the molecular pathogenesis of renal lesions caused by *Txn1*-F54L mutation, we conducted RNA-seq analysis. Figures 5a, 5b, and 5c show the results of RNA-seq of WT and homozygotes for the principal component analysis, heatmap, and MA plots, respectively. The top 10 cellular components obtained from GO enrichment analysis of genes with decreased expression (*P* < 0.05) were mainly associated with the mitochondria, apical plasma membrane, and brush border membrane, which are localized in the paroxysmal tubules (Figure 5e and Supplementary Table S1). Differentially expressed genes (*P* < 0.05) associated with mitochondria were extracted from the gene lists, and their Z-scores are presented as a heatmap (Figure 5d). TEM images of paroxysmal tubules (Figure 5f) in homozygotes revealed a significant decrease in the number of mitochondria (Figure 5g) (*P* < 0.01) and decreased length/ width ratio of mitochondrial (Figure 5h) (*P* < 0.01).

**Figure 5.**
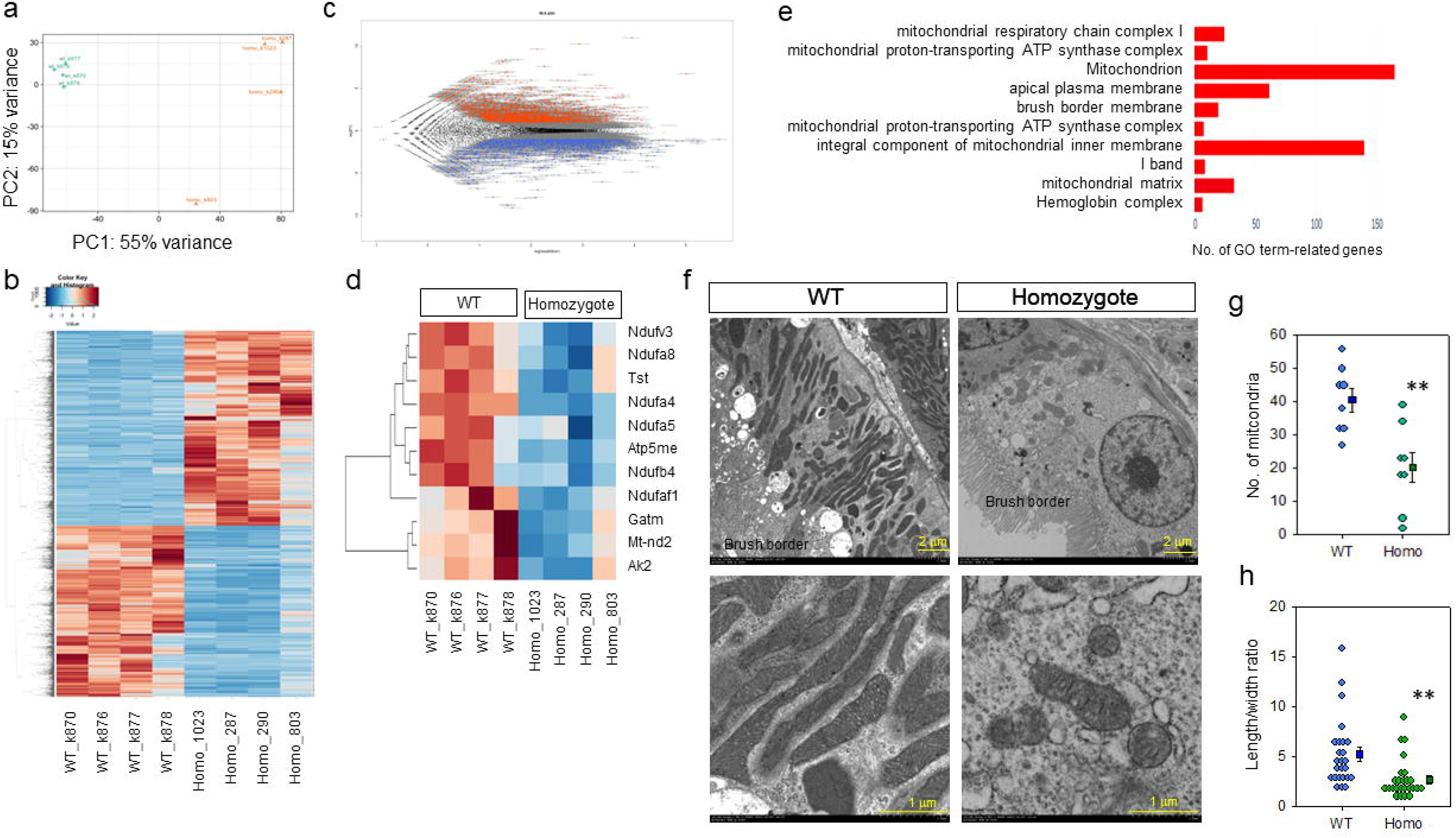
Results of RNA-seq. (**a**) Principal component analysis (PCA) for rats at 15 weeks of age. Principal component 1 (PC1) indicates the component that maximizes the variability of each sample data. PC2 indicates the second most varied component. (Green: WT; Orange: homozygotes). (**b**) Hierarchical clustering analysis and heatmap of differential expression genes. (**c**) MA plot. (**d**) Heatmap of differential expression genes (*P* < 0.05) related to mitochondria. (**e**) The top 10 downregulated cellular components with *P* < 0.05 for GO enrichment analysis. (f) TEM images of paroxysmal tubules and mitochondria. (g) The number of mitochondria / 100 μm2 of TEM image area. (h) The length/width ratio of the mitochondria in the paroxysmal tubules. ** *P* < 0.01

### Significant upregulation of inflammation and regulated cell death pathways

The top 30 genes obtained from the GO enrichment analysis of genes with increased expression (*P* < 0.05) were mainly associated with inflammation, apoptosis, aging, and tissue repair (Figure 6a and Supplementary Table S2). The Z-scores of the genes related to inflammation and the regulated cell death pathways are presented in a heatmap (Figure 6b). The TPM values of pyroptosis-related, apoptosis-related, and necroptosis-related genes are summarized in Supplementary Table S3. IHC was performed to confirm the localization of the upregulated genes using anti-NLRP3, anti-GSDMD, anti-CASP3, and anti-RIPK3 antibodies. They stained strongly in the tubules of homozygotes. Anti-8-OHdG antibody, which indicates DNA damage due to oxidative stress, was also strongly stained in the tubules (Figure 6c).

**Figure 6.**
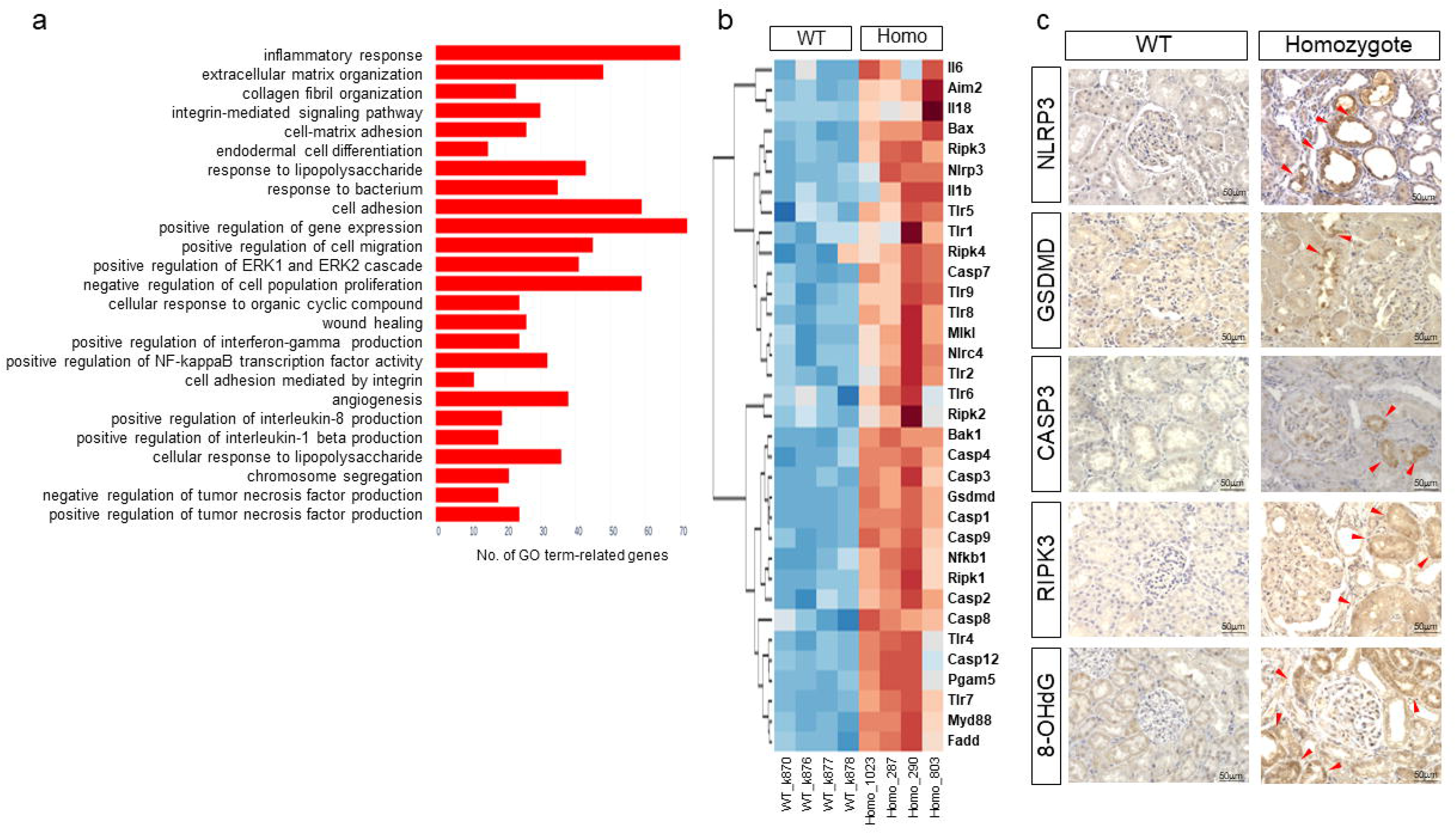
Inflammation and regulated cell-death pathways. (**a**) The top 30 upregulated biological process with *P* < 0.05 for GO enrichment analysis. (**b**) Heatmap of differential expression genes (*P* < 0.05) related to inflammasome and regulated cell-death pathways. Comparison of TPM values of WT vs. homozygotes for pyroptosis (Aim2-Casp1-Nlrc4-Nlrp3-Gsdmd axis), inflammation (Nfkb1-Il1b-Il18 axis), apoptosis (Bak1-Bax-Casp9-Casp3 axis), and necroptosis (Pgam 5-Ripk1-Ripk3-Mlkl axis). (**c**) Immunohistochemistry at end-stage of homozygote (4 months) and WT (4 months). Markers of pyroptosis (NLRP3 and GSDMD), apoptosis (CASP3), necroptosis (RIPK3), and oxidative DNA damage (8-OHdG) were prominent at homozygous tubules. Red arrows indicated strongly stained lesions.

No significant changes were observed in the expression of genes involving *Txn/Txnip/Txnrd/Prdx* system (Supplementary Table S4).

### Confirmation of CKD reproducibility

We performed experiments to confirm the reproducibility of CKD caused by the Txn1-F54L mutation. A short lifespan was also observed in homozygous *Txn1*-F54L rats generated using CRISPR/Cas9 genome editing (Figure 7a). High BUN levels with marked elevation at the end stage (Figure 7b), hypoalbuminemia (Figure 7c), and high T-Cho levels (Figure 7d) were observed; there were no statistical differences in P, Ca, AST, and CK levels (Figure 7e-7h). Histopathology of various organs throughout the body revealed marked nephrosclerosis, thymic regression, and degeneration of the vascular tunica media (Figure 7i). The hearts of homozygotes showed no significant differences from those of WT mice. No notable changes were observed in the liver, pancreas, or adrenal glands (data not shown).

**Figure 7.**
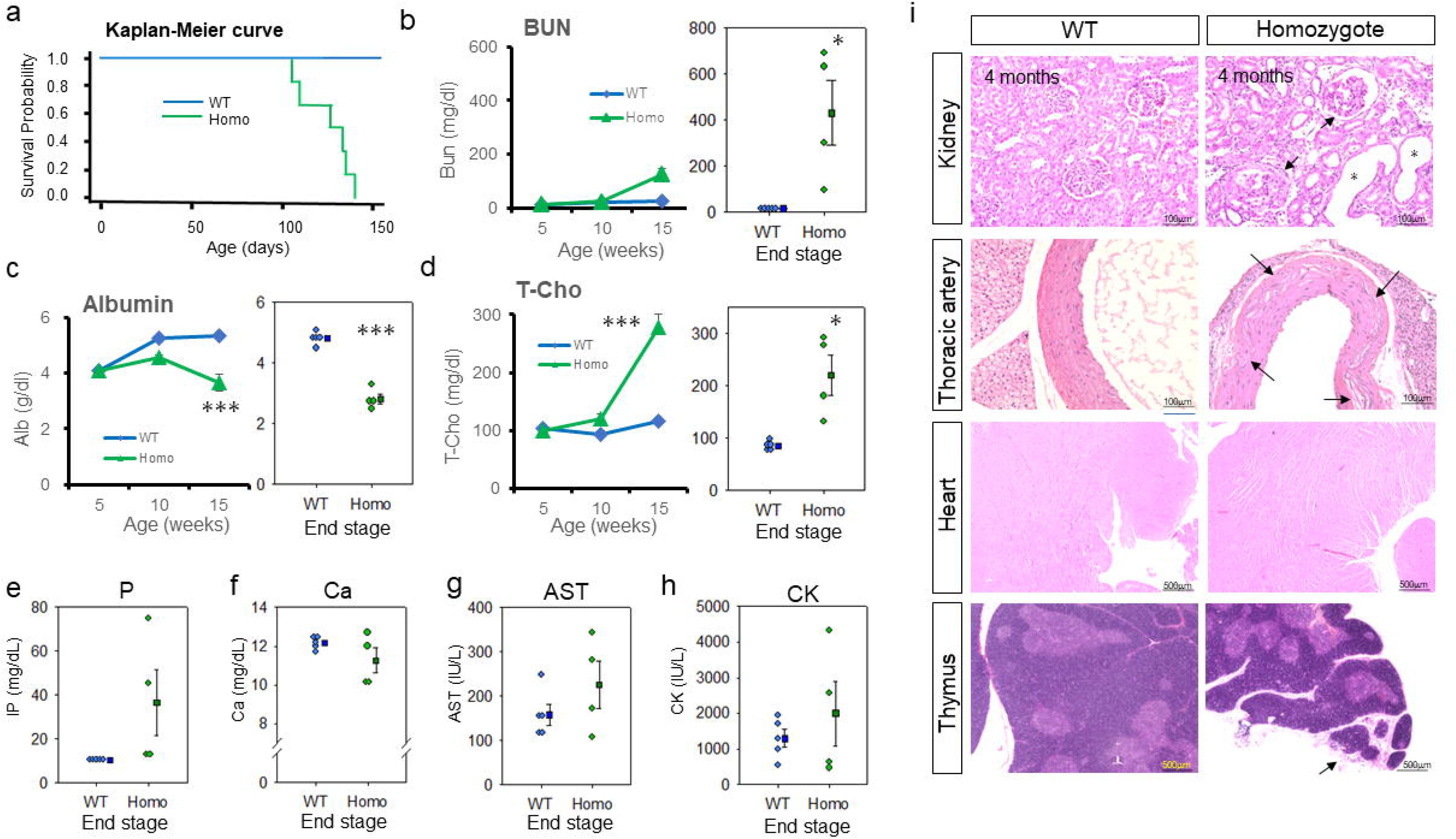
Confirmation of CKD reproducibility. (**a**) Kaplan-Meier survival curves for WT (male: n = 3, female: n = 3) and homozygotes (male: n = 3, female: n = 3). Chemical tests include BUN (**b**), serum albumin (**c**), total cholesterol (**d**), serum phosphate (**e**), serum calcium (**f**), AST (**g**), and CK (**h**). The level of BUN, total cholesterol, and serum albumin were significantly elevated (*P* < 0.05) in homozygous rats. Pathological findings of kidney, thoracic artery, heart, and thymus were presented (**i**).

## DISCUSSION

This study provides a comprehensive understanding of the effects of Txn1 mutations on renal function and sheds light on their underlying pathophysiological basis. First, we showed that a *Txn1* mutation can be a monogenic cause of CKD in rats. Second, CKD is characterized by nephrosclerosis associated with mitochondrial dysfunction, leading to ESKD in all individuals. Oxidative stress markers were prominent in the renal tubules. Third, RNA-Seq analysis showed marked upregulation of genes associated with inflammation and regulated cell death pathways, including apoptosis, pyroptosis, and necroptosis.

### Txn1-F54L is a monogenic cause of CKD

The genetic backgrounds of the rats generated by ENU mutagenesis and the CRISPR/Cas9 system were F344/Nlc and F344/Jcl, respectively. We confirmed that the Txn1-F54L mutation caused CKD in these two strains. CKD progression in F344/Jcl - Txn1-F54L homozygotes was comparable with that in F344/Nlc -Txn1-F54L homozygotes generated via ENU mutagenesis. Our previous study showed that Txn1 is expressed in each organ throughout the body in rats. ^22^ However, kidney was the most prominently affected organ among all the organs as per biochemical and pathological test results. All individuals developed ESKD, leading to premature death. Transgenic mice with Txn1 dominant-negative mutation are susceptible to cardiac hypertrophy induced by transverse aortic constriction. ^24^ Although *Adem* rats also showed myocardial fibrosis and lymphocytic infiltration, these changes were less pronounced and unlikely to cause early mortality. The most notable change, other than in the kidney, was early thymic involution. As T lymphocytes have been reported to decrease in patients with end-stage renal failure, ^25^ thymic involvement in *Adem* rats may have been secondary to CKD. Muscle wasting and uremic inflammation in patients with CKD were reported to lead to premature aging. ^26, 27^ The weight loss, lethargy, and thymic involution in ESKD in *Adem* rats are similar to those observed in humans.

### Phenotype of CKD and underlying pathology in *Adem* rat

Regardless of homozygosity or heterozygosity, the renal pathological findings in all individuals were prominent segmental or global nephrosclerosis, tubular dilation, mesangial cell proliferation, and interstitial fibrosis. Diabetes and hypertension are the major risk factors for CKD. ^28^ The relationship between Txn1/TXNIP and diabetes is well known. However, hyperglycemia was not associated with the onset or progression of CKD in *Adem* rats. In contrast, hypertension and elevated urinary albumin levels developed simultaneously. Systolic and diastolic blood pressure gradually increased during CKD progression. The question arises as to whether hypertension in *Adem* rats is a cause or a consequence of CKD. Animal models of hypertension in mice and rats include angiotensin II-induced hypertension, ^29^ spontaneously hypertensive rats, ^30^ and two-kidney one-clip rats. ^31^ Since the renal lesions in these hypertensive models are mild, ^32^ whereas those in *Adem* rats are severe, hypertension in *Adem* rats may be a consequence of CKD rather than a cause.

### Inflammation and regulated cell death pathways

RNA sequencing revealed that the down-regulated genes were mitochondria. The number of mitochondria in the proximal tubules was significantly reduced in the mutant rats. A marker of oxidative DNA damage for 8-OHdG and inflammasome and cell death markers for NLRP3, GSDMD, CASP3, and RIPK3 were strongly stained in the renal tubules. These results suggest a close link between mitochondrial abnormalities in the proximal tubules and regulated cell death pathways. Reabsorption from the tubules is ATP-dependent, and vast ATP requirements are supplied by the mitochondria. ^33^ Txn1 mRNA and protein are localized in the renal cortex, particularly within the proximal tubules. ^34^ Tubular damage may occur quickly when thioredoxin function is impaired. Mitochondrial abnormalities, inflammation, apoptosis, pyroptosis, and necroptosis are the molecular mechanisms underlying kidney damage in *Adem* rats. Apoptosis is the most studied of the three regulated cell-death pathways. Apoptosis is triggered by changes in mitochondrial outer membrane permeabilization (the intrinsic pathway). Subsequently, the released cytochrome c activates caspases 9/3/7 to induce apoptosis. ^35, 36^ Pyroptosis ^37^ and necroptosis ^38^ are inflammation-related cell death.

Historically, these regulated cell-deaths have been thought to be independent pathways. As research progressed, the crosstalk between mutual pathways became the focus of attention. ^39, 40, 41^ Malireddi et al. have proposed the concept of PANoptosis, which is coexisting pyroptosis, apoptosis, and necroptosis. ^42^ Viral infection, ^43, 44^ bacterial infection, ^45^ retinal ganglion cells in glaucoma, ^46^ and ischemia/reperfusion injury of the brain ^47^ and retinal neurons^48^ have all been linked to PANoptosis. Oxidative stress plays a vital role in the pathogenesis of viral infections ^49^ and ischemia-reperfusion injury. ^50, 51^ Interestingly, *Adem* rats, whose pathogenesis is based on oxidative stress, exhibit upregulation of PANoptosis-related genes. A detailed examination of the molecular mechanisms underlying the pathophysiology in *Adem* rats will lead to the elucidation of the relationship between oxidative stress, mitochondrial dysfunction, and multiple regulated cell-death pathways.

### Application of *Adem* rats for translational research of CKD

The incidence of nephrosclerosis is increasing in aging societies. The pathology of *Adem* rats involves multiple regulatory cell-death pathways. Diabetic nephropathy affects NLRP3-mediated pyroptosis ^52, 53, 54^ and renal fibrosis. ^55^ The Ripk1-meditated necroptosis pathway is involved in renal ischemia/reperfusion injury. ^56^ Renal pathology in *Adem* rats is characterized by interstitial fibrosis and nephrosclerosis, leading to premature death. They share a common molecular basis for nephrosclerosis, diabetic nephropathy, and renal ischemia/reperfusion injury. In addition, they exhibit hypertension, cardiovascular disease, and anemia. Therefore, *Adem* rats could help us understand the pathogenesis of these diseases and develop therapeutic methods to prevent comorbidities.

## AUTHOR CONTRIBUTIONS

I. Ohmori, M. Ouchida, and T. Mashimo designed the study. T. Mashimo generated animals. I. Ohmori and M. Ouchida conducted most experiments, chemical tests, histological assessments, IHC, TEM, and RNA-seq. S. Toyokuni, H. Uchida, and Y. Hada assessed the pathology. I. Ohmori wrote the original draft, and the other authors reviewed and edited the manuscript. All authors have approved the final manuscript.

## ACKNOWLEDGEMENTS

This work was supported by research grants from JSPS KAKENHI (Grant Numbers 16H05354 and 22K07914) and JSPS KAKENHI (grant number JP 16H06276) (AdAMS). We thank Dr. R. Nakaki and Dr. Y. Kondo (Rhelixa Inc.) for analyzing the RNA-seq data. We thank Ms. Yumiko Morishita, Mika Monobe, and Miki Kajino for their technical assistance with histology and IHC. We thank Ms. Masumi Hurutani and Tomomi Tsukano for their technical assistance with TEM. We thank Editage (www.editage.com) for English language editing.

---

**Supplementary figure S1.**
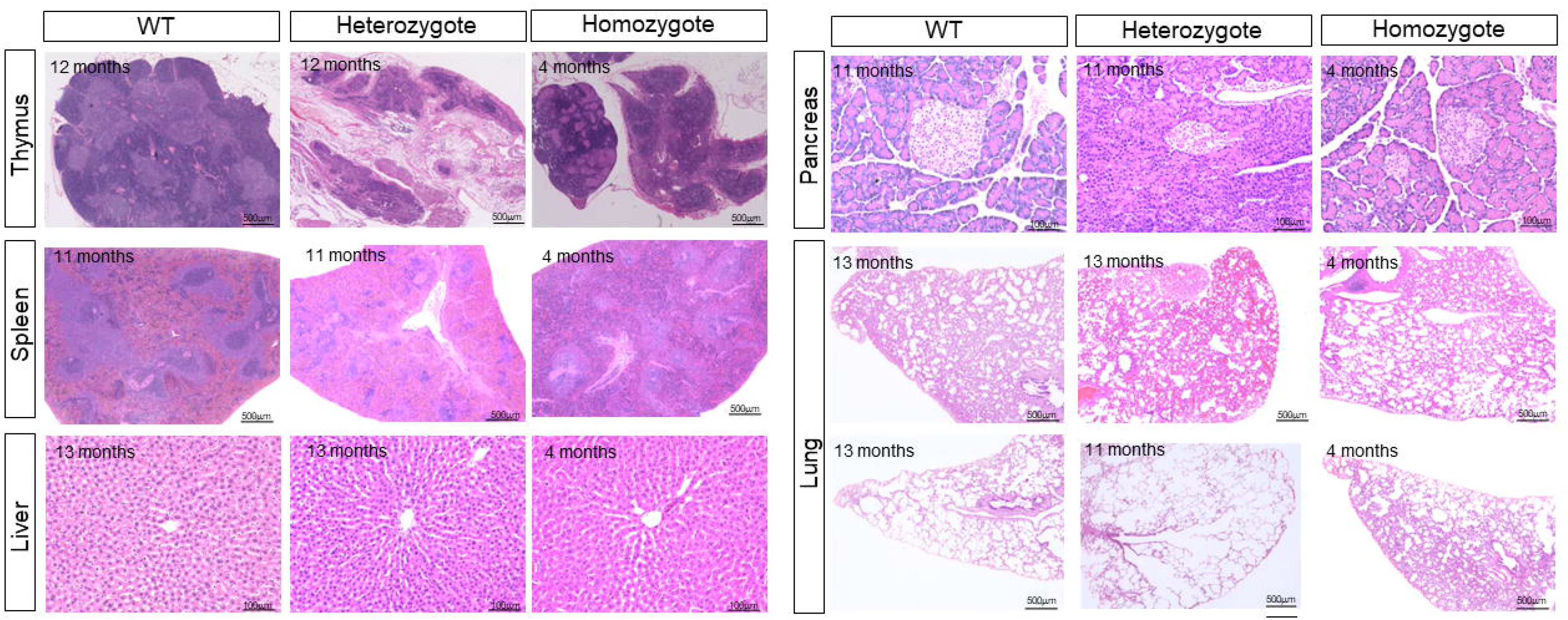
Pathology of other organs. The representative pathological findings of thymus, spleen, liver, pancreas, and lungs.

